# Quantitative Analysis of Plasmodesmata Permeability using Cultured Tobacco BY-2 Cells Entrapped in Microfluidic Chips

**DOI:** 10.1101/2021.02.19.431975

**Authors:** Kazunori Shimizu, Masahiro Kikkawa, Ryo Tabata, Daisuke Kurihara, Ken-ichi Kurotani, Hiroyuki Honda, Michitaka Notaguchi

## Abstract

Plasmodesmata are unique channel structures in plants that link the fluid cytoplasm between adjacent cells. Plants have evolved these microchannels to allow trafficking of nutritious substances as well as signaling molecules for intercellular communication. However, tracking the behavior of plasmodesmata in real time is difficult because they are located inside tissues. Hence, we developed a microfluidic device that traps cultured cells and fixes their positions to allow testing of plasmodesmata permeability. The device has 112 tandemly aligned trap zones in the flow channel. Cells of the tobacco line BY-2 were cultured for 7 days and filtered using a sieve and a cell strainer before use to isolate short cell clusters consisting of only a few cells. The isolated cells were introduced into the flow channel, resulting in entrapment of cell clusters at 25 out of 112 trap zones (22.3%). Plasmodesmata permeability was tested from 1 to 4 days after trapping the cells. During this period, the cell numbers increased through cell division. Fluorescence recovery after photobleaching experiments using a transgenic marker line expressing nuclear-localized H2B-GFP demonstrated that cell-to-cell movement of H2B-GFP protein occurred within 200 min of photobleaching. The transport of H2B-GFP protein was not observed when sodium chloride, a compound known to cause plasmodesmata closure, was present in the microfluid channel. Thus, this microfluidic device and one-dimensional plant cell samples allowed us to observe plasmodesmata behavior in real time under controllable conditions.

## Introduction

In plants, there are microscopic tunnels that penetrate the cell wall and directly connect the cytoplasm of neighboring cells. These tunnels, called plasmodesmata (PDs), are ultrafine structures, often as small as 30 nm in diameter and 100 nm length, and facilitate the movement of materials between cells (Brunkard et al., 2015). It is estimated that about 1,000–100,000 PDs are located between neighboring cells (Maule, 2008; Lucas et al., 2009; Xu and Jackson, 2010). Recent studies have shown that PDs are involved in the transport of proteins, mRNAs, plant hormones, and other substances involved in plant functions. They also allow the transfer of pathogenic bacteria and viruses between cells (Lucas et al., 1995; Turgeon and Wolf, 2009; Lee and Lu, 2011; Brunkard et al., 2015; Han and Kim, 2016). Since PDs are fundamental functional structures for plants and have unique characteristics, they have attracted much attention for decades, and many studies have focused on elucidating their structures and functions in detail.

Because PDs are ultrafine structures with diameters of only several tens nm and are embedded in thick cell walls, the methods for analyzing them are limited. For more than 100 years, it has been extremely difficult to study their functions and structures (Roberts and Oparka, 2003). However, with the development of electron microscopy, research on PDs has made remarkable progress since the first structural details were revealed by Ding and his colleagues in 1992 (Ding et al., 1992). Currently, electron microscopy and confocal microscopy have become the main methods used in research on PDs. These techniques have made it possible to correlate changes in the function of PDs with changes in their structure. For example, it has been reported that differences in the permeability of substances at different growth stages of plants can be related to differences in the structure of PDs (Nicolas et al., 2017), and changes in the permeability of auxin-responsive substances can be related to structural changes around PDs (Han et al., 2014). Genetic screening has also succeeded to identify components regulating PD formation and permeability (Stonebloom et al., 2009; Xu et al., 2012; Benitez-Alfonso et al., 2013; Brault et al., 2019).

However, there are still three major issues with these research methods. First, confocal microscopy using fluorescent proteins as reporters has a spatial resolution of 250–100 nm, which is insufficient for detailed observations of PDs. Second, it takes a long time to prepare samples for microscopic observation, and it often takes about 2 to 4 weeks to grow a plant for observation. Third, in the case of electron microscopy, it is necessary to fix the tissue for observation, so it is impossible to study PD behavior in real time in live tissues. In addition, fixation and preparation of sections require skills, making it difficult to perform simple experiments.

The cultured cells used in this study are expected to be useful for analyses of PDs because the culture cells position in one dimensional allowing us to clearly observe. In fact, intercellular localization of PD localized proteins has been studied using cultured cells (Knox et al., 2015). However, floating cultured cells are difficult to observe over time. Moreover, fixing the position of the object is important especially for high-resolution imaging. Therefore, to further understand PD behavior, a simple *in vitro* PD analysis method using cultured cells that can be observed over time is necessary.

In this study, we aimed to establish a method to assess the permeability of PDs using a microfluidic device in which cultured plant cells can be fixed in position. We used the tobacco (*Nicotiana tabacum*) BY-2 cell line, which has been used widely as a model cell in plant cell biology. The doubling time of BY-2 cells is 12–14 h, which is as short as that of plant cells (Geelen and Inze, 2001; Miyazawa et al., 2003). In addition, BY-2 cells form small clusters consisting of several cylindrical cells connected in tandem in liquid culture medium. This characteristic makes it possible to observe permeability via PDs between adjacent cells under a microscope. First, we observed the properties of BY-2 cells, including their length and linearity, during culture to select the appropriate duration of culture for cells used in this study. Next, to avoid changing the position of BY-2 cells during the experiments, we fabricated a microfluidic device with a microchannel to trap the BY-2 cells. Finally, we explored the permeability of PDs in trapped BY-2 cells by quantitatively assessing fluorescence recovery after photobleaching (FRAP).

## Materials and Methods

### Culturing of tobacco BY-2 cells

Tobacco BY-2 cells were cultured in modified Linsmaier and Skoog (LS) medium (see the following recipe, Nagata et al., 1992). The BY-2 cell cultures were maintained at 26°C with shaking at 130 rpm in the dark on a shaking stirrer (NR-3, Titech Co., Ltd., Aichi, Japan) in an incubator (IS-2000, Toyo Seisakusho Co., Ltd., Chiba, Japan). The medium was dispensed into 300-mL flasks (95 mL medium/flask). Each flask was capped with a silicon stopper, and autoclaved at 120°C for 20 minutes. For subculture, 3 mL cell suspension was removed from the flask on the 7^th^ day of culture and transferred to a new flask. For incubation of the H2B-GFP strain, 50 mg/L kanamycin sulfate (117-00341, Fujifilm Wako Pure Chemicals Co., Ltd., Osaka, Japan) was added to the culture medium.

Recipe for modified LS medium

- Murashige and Skoog plant salt mixture (392-00591, Fujifilm Wako), 4.6 g/L
- Sucrose (193-00025, Fujifilm Wako), 30 g/L
- Potassium Dihydrogen Phosphate (169-04245, Fujifilm Wako), 0.2 g/L
- Myo-inositol (I5125, Sigma, St. Louis, MO, USA), 0.1 g/L
- Thiamin hydrochloride (205-00855, Fujifilm Wako), 1.0 mg/L
- 2.4-dichlorophenoxyacetic acid (040-18532, Fujifilm Wako), 0.2 mg/L

Adjust pH to 5.8 with KOH

### Plasmid construction and transformation of tobacco BY-2 cells

To construct HSpro::H2B-sGFP (designated as DKv813), the 446-bp upstream region (−506 to −61 bp) of soybean *Gmhsp17.3-B* (Schöffl et al., 1984) and the full-length coding region of *H2B* (HTB1:At1g07790) fused to sGFP were cloned into the binary vector pPZP211 (Hajdukiewicz et al., 1994). To generate transgenic BY-2 cell lines expressing H2B-GFP, *Agrobacterium-mediated* transformation was performed as previously described (An, 1985). Transformants were selected on modified LS medium containing 1.5% (w/v) agar and 50 μg mL^-1^ kanamycin and then cultured for 3 weeks before initiating a liquid culture in modified LS medium. The callus cells were transferred to liquid medium using a sterilized platinum loop. The cells were subcultured once or twice before use to stabilize growth. The HSpro::H2B-sGFP transgenic cell line constitutively expresses H2B-sGFP without heat shock treatment in medium culture.

### Microdevice fabrication

The microfluidic devices were fabricated by replica molding using polydimethylsiloxane (PDMS; SILPOT 184, DuPont Toray Specialty Materials K.K., Tokyo, Japan) as we reported previously, with some modifications (Yamaoka et al., 2019). Briefly, the silicon wafer was spin-coated with SU-8 3050 (MicroChem Corp., Newton, MA, USA) at 1750 rpm for 60 s and baked at 95°C for 30 min. The wafer was exposed to ultraviolet light through a photomask and baked at 95°C for 10 min. After development using SU-8 developer (MicroChem Corp.), PDMS mixed with a curing reagent at a ratio of 10:1 was poured onto the mold, and then cured by baking at 70°C for 60 min. Two 1.5-mm diameter holes were made for an inlet and outlet using a biopsy needle (Kai Industry, Gifu, Japan). Then, the PDMS and a cover glass were treated with air plasma and bonded to each other.

### Preparation and image analysis of BY-2 cells

To prepare small clusters of BY-2 cells, the BY-2 cell culture was passed through a 140-μm sieve (NRK, Tokyo, Japan) and subsequently through 100, 85, 70, 50, and 40-μm cell strainers (pluriSelect Life Science, Leipzig, Germany). Before and after separation, a 1-mL aliquot of BY-2 cell culture was mixed with CellTracker Green CMFDA Dye (C7025, Thermo Fisher Scientific, Tokyo, Japan) at 5 μM and incubated for 30 min at 26°C with agitation. Then, 10 μL BY-2 cell-containing medium was placed on the slide glass and covered with a cover glass. Fluorescence images were obtained using a fluorescence microscope (BZ-X700, Keyence, Osaka, Japan). The images were analyzed by ImageJ; the objects were recognized by color threshold and subsequently the objects crossing the edges were excluded. Then, the minimum length of the minor axis of the approximate minimum ellipse and area of the objects, *i.e*., BY-2 cell clusters, were measured. These procedures are illustrated in Figure S1.

### Introducing BY-2 cells into the microfluidic device

The device was sterilized by exposure to ultraviolet light for 1 h and treated with air plasma to make the surface of the microchannels hydrophilic. Then, 700 μL medium without cells was placed into the inlet and allowed to fill the channels. The cell suspension was prepared at a concentration of 500–1000 cells/mL. A 700-μL aliquot of the cell suspension was introduced slowly into the channels from the inlet. Then, a 700-μL aliquot of medium without cells was introduced to remove the un-trapped cells.

### Culture of trapped BY-2 cells in the microfluidic device

After introducing cells into the channels, the inlet and outlet of the microchannels were plugged with P200 filter pipette tips (127-200XS, Watson, Tokyo, Japan) filled with/without culture medium to prevent the channels from drying. Then, the device with tips was incubated at 26°C in the dark. The medium was exchanged every 24 h by introducing 700 μL medium slowly from the inlet.

### Quantitative measurement of PD permeability

A culture of BY-2 cells expressing H2B-GFP was prepared at 500–1000 cells/mL as described above. To avoid unintended physical stress during the cell trapping procedure that may close PDs, and to increase the number of clusters with three or more cells, we cultured the trapped BY-2 cells for 1 day or longer.

A confocal microscope (LSM510/LSM5 Pascal, Zeiss, Germany) equipped with 488 nm argon laser (LGK7872ML, LASOS Lasertechnik GmbH, Jena, Germany) was used for observation and photobleaching. For photobleaching, the settings were as follows: objective lens, Plan-Neofluor 10x/0.3; pinhole, max (141.4 μm); laser power, 100%; observation area, 898.2 μm × 898.2 μm; laser irradiation area, 219.1 μm × 219.1 μm; photobleaching period by laser scanning, 963.04 msec; irradiation duration, 25 min. We obtained fluorescence images of the trapped targeted cluster three times (before, just after, and 200 min after photobleaching). Image analysis was performed using ImageJ. The objects and nuclei were identified and their mean fluorescence intensity was compared.

### Statistical analysis

Statistical analysis was performed using Tukey’s HSD for independent samples with unequal variance using Python version 3.7.4 and its library modules including NumPy (1.17.2), Pandas (0.25.1), and statsmodels (0.12.1). A p-value less than 0.05 was considered statistically significant.

## Results and Discussion

### Analysis of shapes of BY-2 cells during culture

Since BY-2 cells form short cell clusters consisting of several cylindrical cells connected in tandem in liquid culture medium (Figure 1A), we anticipated that it should be relatively easy to observe the exchange of molecules between cells via PDs. Thus, to design a microfluidic device that can fix the position of BY-2 cells, we observed the shapes of BY-2 cells under a microscope.

**Figure 1.**
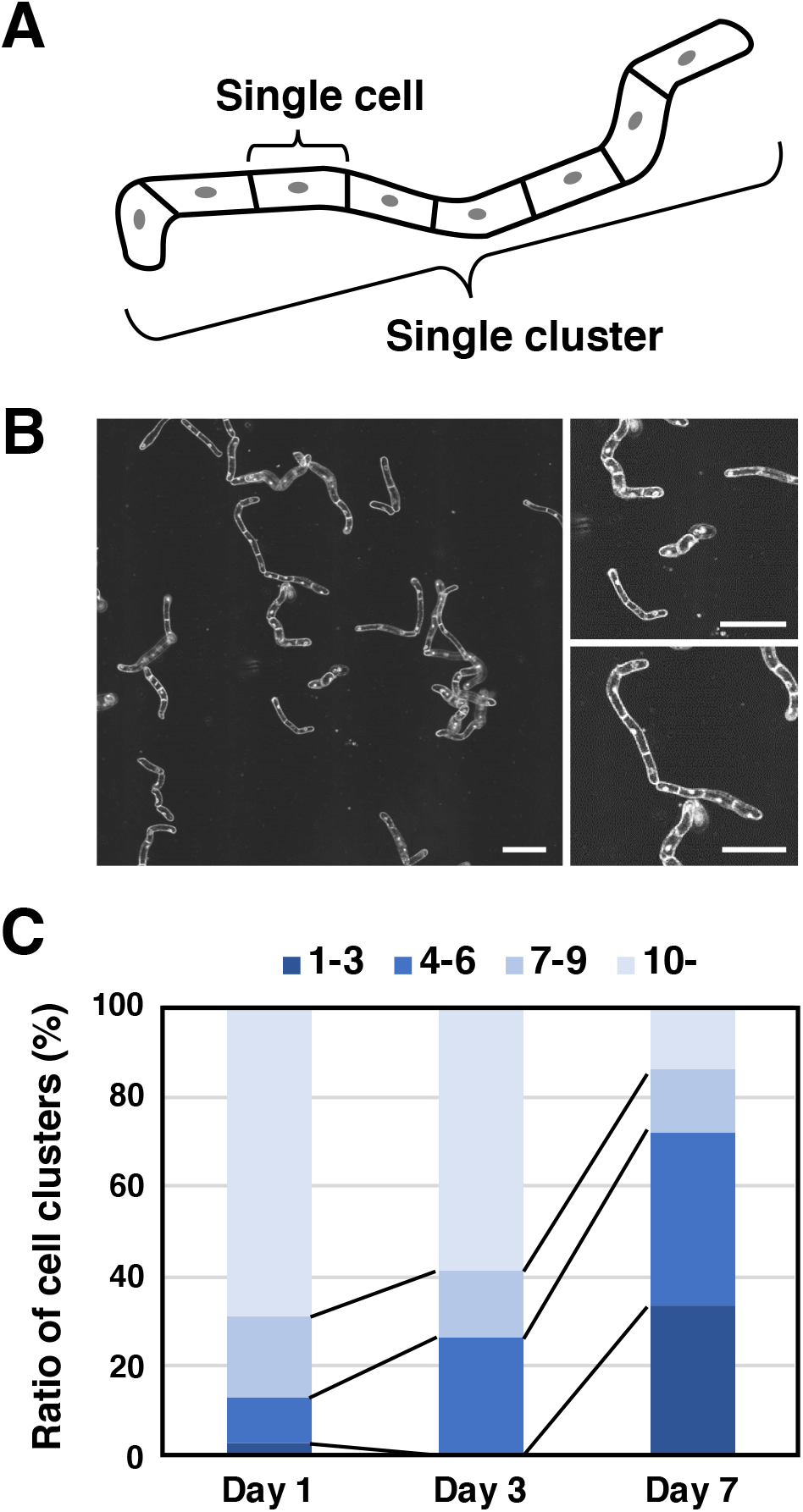
Properties and shapes of BY-2 cells. **(A)** Drawing of a BY-2 cell. **(B)** Representative images of BY-2 cells under microscope. Scale bars, 200 μm. **(C)** Distribution of number of cells in a single cluster of BY-2 cells on days 3, 5, and 7 of culture.

Our observations revealed that the BY-2 cells were linear but not uniform in shape and size. In particular, the degree of bending was different in each cluster (Figure 1B). Since BY-2 cells cultured in flask batches were passaged every 7 days, we counted the number of cells in one cluster on days 3, 5, and 7. The percentages of clusters containing 10 or more cells were 69.2% (27/39) on day 3, 58.8% (20/34) on day 5, and only 14.0% (8/57) on day 7. The percentage of clusters containing four or fewer cells increased to 64.9% (37/57) on day 7 in our conditions (Figure 1C). Since the mitotic index usually peaked at around days 2 and 3 and declined rapidly (e.g. Magyar et al., 2005), the prolonged cell culture period might cause break down the long cell cluster to a short.

### Design and fabrication of microfluidic device for fixing the position of BY-2 cells

We designed and fabricated a microfluidic device for fixing the position of BY-2 cells referring to Kim et al. 2014 (Figure 2A). In the previous report, cell sorting and trapping were demonstrated using polystyrene microspheres (sizes: 15 μm, 6 μm, and 4 μm) and three different waterborne parasites including *Giardia* cysts (ellipsoid with short and long axes of 8 μm and 19 μm, respectively). To trap BY-2 cells, in this study, we modified the dimensions and the shape of the trap zones in the microfluidic device. The device has a main channel and a side channel that are connected by a trap zone (Figure 2B). The cell suspension was added to the main channel through the inlet. According to the simulation by Kim et al. 2014, the fluidic pressure of the main channel is always higher than that of the side channel at the same position because the ratio of the width of the inlet and outlet of the trap zone, the main channel, and the side channel is set to 2:1:8:20. In our preliminary tests, the fluid flowed from the main channel to the side channel through the trap zone without backflow, and the cells were trapped in the trap zone.

**Figure 2.**
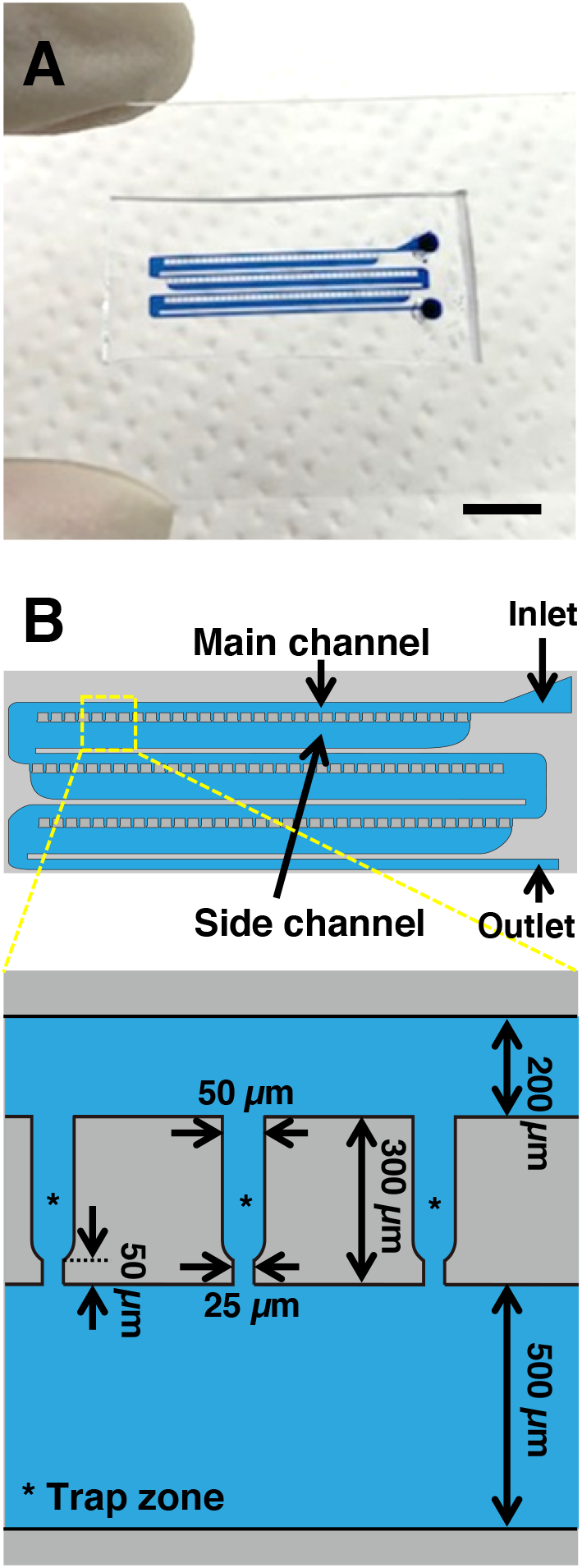
Design and fabrication of microfluidic device. **(A)** Fabricated microdevice with channels filled with blue-colored water. Scale bar, 5 mm. **(B)** Shape and size of microchannels.

In this study, we designed each trap zone to entrap one short cluster of BY-2 cells (Figure 1C). The entrance of the trap zone was 50 μm, which is almost the same width of a single cluster of BY-2 cells. The exit of the trap zone was 25 μm, which is narrower than the cell width. The length of the trap zone was set to 300 μm, which is about the length of a cluster of several cells. The device had a total of 112 trap zones (Figure 2B).

### Introducing BY-2 cells into the microfluidic device

We examined whether short clusters of BY-2 cells could be trapped in the developed device. To clearly observe the location of cells in the device, the cells were stained by CellTracker Green before use. We introduced an aliquot of cells cultured for 7 days, which contained many small cell clusters, into the inlet (Figure 1C). However, the cells became clogged in the main channel near the inlet and were not trapped in the trap zone. Microscopic observations revealed that this was caused by clogging of the main channel with long cell clusters in the cell suspension. To remove the long clusters, the cell culture was passed through a sieve and a series of cell strainers (see Materials and Methods). The cell suspension after this separation step contained shorter and straighter cell clusters (Figure 3A). When the separated cells were introduced into the device, the cells in short clusters were trapped in the trap zone (Figure 3B). On average, 25 ± 4.5 cell clusters were trapped by the device with 112 trap zones (entrapment frequency, 22.3% ± 4.1%). The number of cells in the trapped cluster at the trap zone was counted. More than half of the clusters consisted of two cells, the other clusters consisted of one cell or three or more cells (Figure 3C). Thus, the short clusters were successfully entrapped by the microfluidic devices.

**Figure 3.**
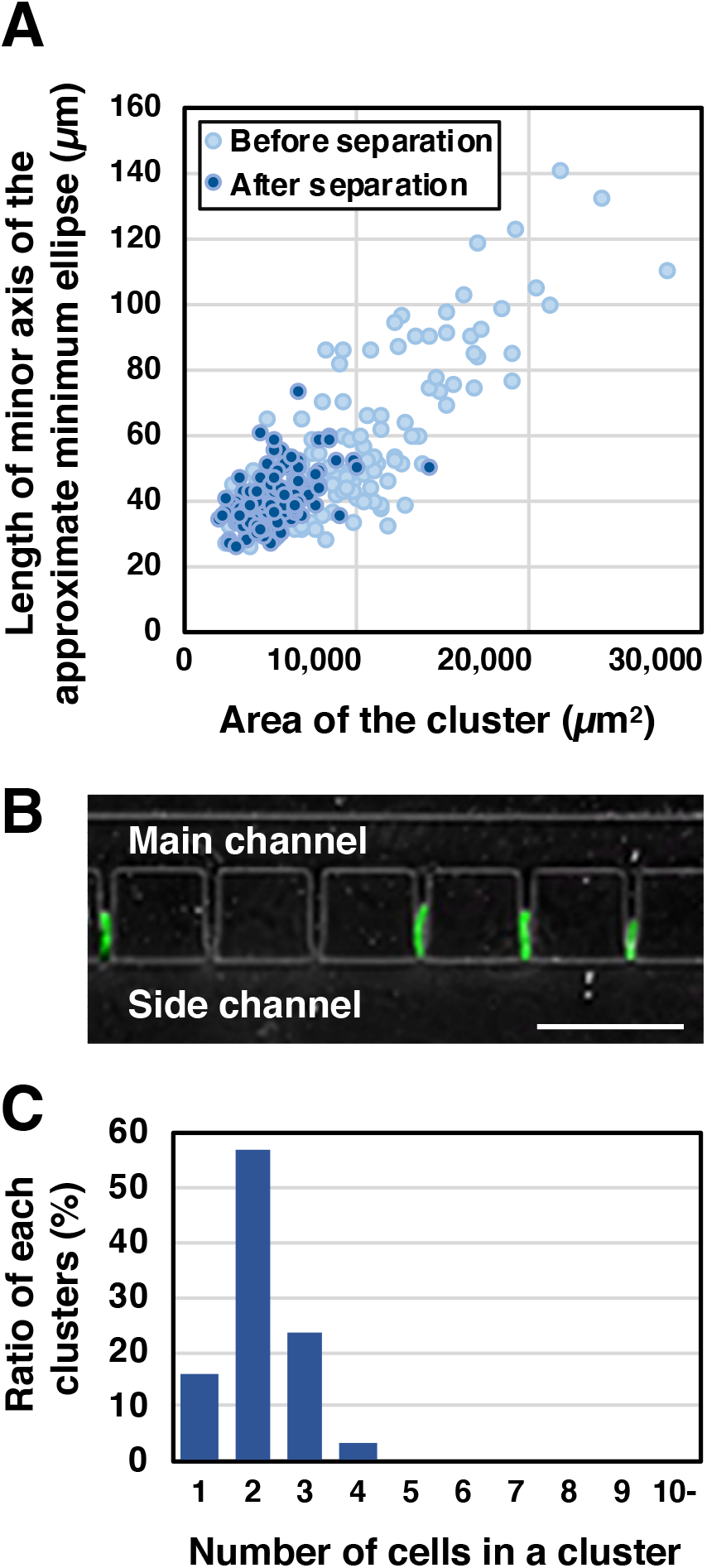
Trapping of BY-2 cells in the microfluidic device. **(A)** Relationship between length of minor axis of approximate minimum ellipse and area of cell cluster. Closed circle: before cell separation. Open circle: after cell separation. BY-2 cells stained with CellTracker Green were used and 150 clusters were analyzed for each condition. **(B)** Representative image of trapped cells. Scale bar, 500 μm. **(C)** Ratio of number of cells in the trapped clusters in microfluidic devices.

### Cultivation of trapped BY-2 in the microfluidic device

To investigate PD permeability using the trapped BY-2 cells, it was important that the cells remained undamaged. We investigated whether the trapped BY-2 cells retained their proliferation ability in the device. As shown in Figure 4A, the number of cells in the trapped single cluster at the trap zone increased over time through cell division and the cluster became longer. The average number of cells in one trapped cluster was 1.6 ± 0.5 immediately after trapping and increased to 13.6 ± 3.8 after 4 days of culture (Figure 4B). These results confirmed that trapping by this device did not inhibit cell proliferation.

**Figure 4.**
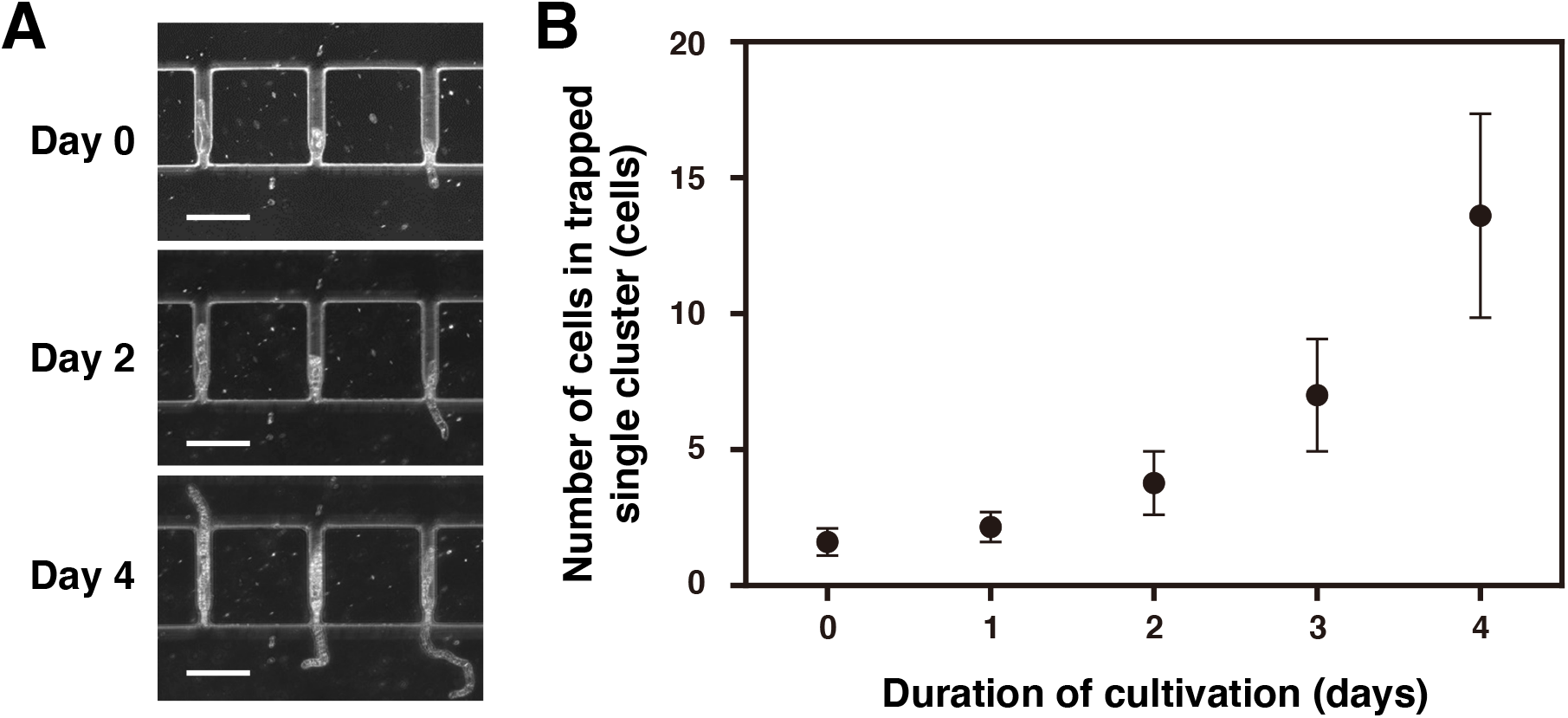
Culturing of trapped BY-2 cells inside the microfluidic device. **(A)** Representative image of trapped cells on days 0, 2 and 4 after trapping. Scale bars, 200 μm. **(B)** Change in number of cells in a trapped single cluster (13 clusters were analyzed).

We calculated the percentage of cells that remained trapped until day 4 (number of clusters trapped at the trap zone on day 0 / number of clusters trapped at the trap zone on day 4 × 100). This analysis revealed that 61.6% ± 7.2% of cells remained trapped on day 4. It is likely that the medium flow released some of the trapped cells from the trap zone during 4 days of culture. In future studies, it should be possible to adjust the trap zone shape and precisely control the flow velocity of the medium to increase the percentage of trapped cells.

### Quantitative evaluation of PD permeability by FRAP

We used the BY-2 cells trapped in the trap zone in the microfluidic device to quantitatively evaluate PD permeability using FRAP. The cytoplasm of each cell in a single cluster is separated by PDs and molecules in the cytoplasm move between the cells via the PDs. In this study, we used BY-2 cells expressing a fusion protein of histone H2B and GFP (H2B-GFP). In each cell, H2B-GFP was translated in the cytoplasm and translocated to the nucleus (Figure S2A). We expected that some H2B-GFP would permeate through the PDs and diffuse into the cytoplasm of adjacent cells, and then be translocated to the nucleus as observed for the nuclear localization signal fused GFP and endogenous transcription factors such as KNOTTED1, LEAFY and SHORTROOT (Jackson et al., 1994; Sessions et al., 2000; Nakajima et al., 2001; Wu et al., 2003). In our experimental environment, when we photobleached cells of a part of the cluster, the GFP fluorescence of photobleached cells recovered by about 140 minutes after laser irradiation. However, when all the cells in the cluster were irradiated by the laser, no recovery of fluorescence was observed by 140 minutes, suggesting that the recovery of fluorescence was not due to accumulation of GFP produced via *de novo* protein expression, but rather to GFP that migrated from neighboring cells through PDs (Figures S2B,C). Consequently, we decided to evaluate the recovery of GFP fluorescence at 200 min after laser irradiation. In the experiment, we used cell clusters consisting of three or four cells (Figure 5A). As the negative control (NC), all cells in the cluster, including the cell at position 0, were irradiated with the laser and photobleached. In the experimental treatment, the cell at position 0 was not laser irradiated, while those at positions 1 and 2 at the end of the cell cluster were irradiated with the laser and photobleached selectively. At 200 min after photobleaching, the recovery of fluorescence in the cells at positions 1 and 2 was observed and quantified. By comparing the fluorescence intensity between NC cells and those in the experimental treatment, the movement of H2B-GFP from the cells at position 0 to those at positions 1 and 2 was assessed.

**Figure 5.**
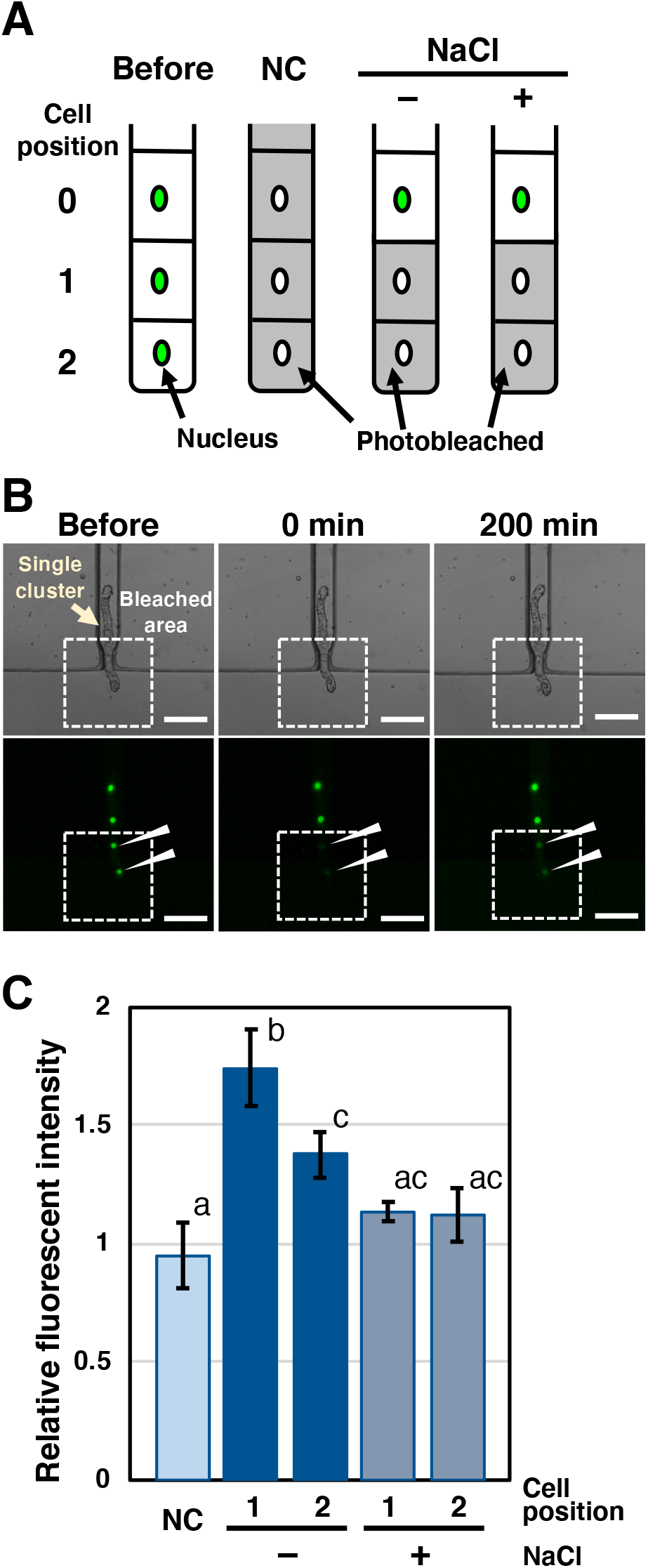
Quantitative analysis of plasmodesmata permeability. **(A)** Schematic illustration of conditions used in this experiment. NC, negative control. **(B)** Representative images of fluorescence recovery after photobleaching (FRAP) in experimental conditions (without NaCl). Slashed rectangles indicate laser-irradiated areas. Fluorescence intensities of photobleached nuclei were recovered by 200 min after laser exposure (arrowheads). Scale bars, 100 μm. **(C)** Quantitative comparison of relative fluorescence intensity. Results of Tukey’s HSD test are shown, *p* < 0.05, *n* = 12, 4, and 5 for negative control (NC) and experimental conditions without and with 100 mM NaCl, respectively.

Figure 5B shows the FRAP results obtained under these experimental conditions. At 200 min after photobleaching, the fluorescence of the photobleached nuclei was recovered, whereas no recovery was observed in the NC cells. In the experimental treatment and the control, recovery to the original fluorescence level was observed at 24 h after photobleaching (Figure S2D). These results suggest that the recovery of fluorescence was caused by the diffusion of H2B-GFP from the cell at position 0 via PDs. In other studies, non-specific trafficking of GFP protein in tobacco and Arabidopsis leaves was also observed in a similar time frame, from several tens of minutes to several hours (Oparka et al., 1999; Kawade et al., 2013). Therefore, BY-2 cultured cells appear to have functional PDs like those in intact plant tissues.

The FRAP results are shown in Figure 5C. The fluorescence intensity of cells at position 1 and 2 was significantly higher than that of NC cells. Furthermore, the fluorescence intensity of cells at position 1 was higher than that of cells at position 2, suggesting that the H2B-GFP produced in non-bleached cells in position 0 was transported to adjacent cells in sequence via PDs. Considering these results, we can conclude that PD permeability was successfully quantified by this FRAP-based method using BY-2 cells trapped in a microfluidic device.

Finally, we performed a similar experiment using trapped cell clusters treated with 100 mM sodium chloride (NaCl). In another study, treatment of Arabidopsis leaves with 100 mM NaCl imposed osmotic stress on cells, resulting in PD closure and a reduction in molecular permeability (Grison et al., 2019). However, it was unknown whether treatment with 100 mM NaCl would reduce the permeability of PDs in BY-2 cells. We found that the recovery of fluorescence intensity was significantly suppressed by treatment with 100 mM NaCl (Figure 5C). Thus, treatment with 100 mM NaCl decreased PD permeability in BY-2 cells. These results provided further evidence that PD permeability could be successfully quantified in entrapped cultured BY-2 cells.

## Conclusion

We have developed a microfluidic device to trap short clusters of BY-2 cells. By using trapped BY-2 cells expressing H2B-GFP and a FRAP technique, we quantitatively evaluated the permeability of PDs. This technology will be useful to test the effect of various compounds on PD permeability. Furthermore, combining this method with mutant strains and/or super-resolution imaging technologies with marker proteins will allow for further in-depth studies on the molecular mechanisms underlying the function and regulation of PDs, and on changes in their transport capacity over time.

## Conflict of Interest

The authors declare that the research was conducted in the absence of any commercial or financial relationships that could be construed as a potential conflict of interest.

## Author Contributions

KS, RT, HH, and MN conceived this study. MK designed and conducted the main experiments with advice from KS, RT, HH, and MN. DK constructed transgenic BY-2 cell lines. MK and RT maintained and prepared the BY-2 cells for experiments. MK and KK performed FRAP experiments. KS, MK, RT, KK, and MN wrote the paper.

## Funding

This work was supported by grants from the Japan Society for the Promotion of Science Grants-in-Aid for Scientific Research (18K05373 and 20H05501 to RT, 18KT0040, 18H03950, and 19H05361 to MN and 20H03273 to KK and MN), Grant-in-Aid for Scientific Research on Innovative Areas (20H05358 to DK), the Japan Science and Technology Agency PRESTO program (JPMJPR18K4 to DK), and the Canon Foundation (R17-0070 to MN).

## Acknowledgments

We thank Kazuki Yakamoto, Haruo Kassai, and Ikue Yoshikawa for technical assistance.

---

**Supplementary Figure 1** Procedure for image analysis of BY-2 cells using ImageJ.

**Supplementary Figure 2** Determination of fluorescence recovery after photobleaching (FRAP) experimental conditions. **(A)** Laser scanning microscope image of BY-2 cells expressing H2B-GFP. Slashed rectangle indicates laser-irradiated area in (B). **(B)** Representative images of FRAP. Yellow and white arrowheads indicate nuclei with recovered and unrecovered GFP fluorescence, respectively. Slashed rectangles indicate areas magnified in (C). **(C)** Magnified fluorescence image of recovered GFP. **(D)** Microscopic images of BY-2 cells in microfluidic device before and after photobleaching. Slashed rectangles indicate areas of fluorescence images. Circles indicate signal from *de novo*-expressed GFP at 24 h after photobleaching. Confocal laser microscopy images were merged from 10 consecutive optical sections. Fluorescence images were merged with bright-field images in (A, B). Scale bars, 200 μm (A–C), 50 μm (D).

## Notes

### Competing Interest Statement

The authors have declared no competing interest.

## References

An, G.H. (1985). High-efficiency transformation of cultured tobacco cells. Plant Physiology 79, 568–570. doi:10.1104/pp.79.2.568

Benitez-Alfonso, Y., Faulkner, C., Pendle, A., Miyashima, S., Helariutta, Y., and Maule, A. (2013). Symplastic intercellular connectivity regulates lateral root patterning. Dev. Cell 26, 136–47. doi: 10.1016/j.devcel.2013.06.010

Brault, M. L., Petit, J. D., Immel, F., Nicolas, W. J., Glavier, M., Brocard, L., et al. (2019). Multiple C2 domains and transmembrane region proteins (MCTPs) tether membranes at plasmodesmata. EMBO Rep. 20, e47182. doi: 10.15252/embr.201847182

Brunkard, J.O., Runkel, A.M., and Zambryski, P.C. (2015). The cytosol must flow: intercellular transport through plasmodesmata. Current Opinion in Cell Biology 35, 13–20. doi:10.1016/j.ceb.2015.03.003

Ding, B., Turgeon, R., and Parthasarathy, M.V. (1992). Substructure of Freeze-Substituted Plasmodesmata. Protoplasma 169, 28–41. doi:10.1007/Bf01343367

Jackson, D., Veit, B., Hake, S.. (1994). Expression of maize *KNOTTED1* related homeobox genes in the shoot apical meristem predicts patterns of morphogenesis in the vegetative shoot. Development 120, 405–413.

Geelen, D.N.V., and Inze, D.G. (2001). A bright future for the bright yellow-2 cell culture. Plant Physiology 127, 1375–1379. doi: 10.1104/pp.010708

Grison, M.S., Kirk, P., Brault, M.L., Wu, X.N., Schulze, W.X., Benitez-Alfonso, Y., Immel, F., and Bayer, E.M. (2019). Plasma Membrane-Associated Receptor-like Kinases Relocalize to Plasmodesmata in Response to Osmotic Stress. Plant Physiology 181, 142–160. doi: 10.1104/pp.19.00473

Hajdukiewicz, P., Svab, Z., and Maliga, P. (1994). The small, versatile pPZP family of Agrobacterium binary vectors for plant transformation. Plant Molecular Biology 25, 989–994. doi: 10.1007/BF00014672.

Han, X., Hyun, T.K., Zhang, M.H., Kumar, R., Koh, E.J., Kang, B.H., Lucas, W.J., and Kim, J.Y. (2014). Auxin-Callose-Mediated Plasmodesmal Gating Is Essential for Tropic Auxin Gradient Formation and Signaling. Developmental Cell 28, 132–146. doi:10.1016/j.devcel.2013.12.008

Han, X., and Kim, J.Y. (2016). Integrating Hormone-and Micromolecule-Mediated Signaling with Plasmodesmal Communication. Molecular Plant 9, 46–56. doi:10.1016/j.molp.2015.08.015

Kawade, K., Tanimoto, H., Horiguchi, G., and Tsukaya, H. (2017). Spatially Different Tissue-Scale Diffusivity Shapes ANGUSTIFOLIA3 Gradient in Growing Leaves. Biophys. J. 113, 1109–1120. doi: 10.1016/j.bpj.2017.06.072

Kim, J., Erath, J., Rodriguez, A., and Yang, C. (2014). A high-efficiency microfluidic device for size-selective trapping and sorting. Lab Chip 14, 2480–2490. doi:10.1039/c4lc00219a

Knox, K., Wang, P., Kriechbaumer, V., Tilsner, J., Frigerio, L., Sparkes, I., et al. (2015). Putting the Squeeze on Plasmodesmata: A Role for Reticulons in Primary Plasmodesmata Formation. Plant Physiol. 168, 1563–72. doi: 10.1104/pp.15.00668

Lee, J.Y., and Lu, H. (2011). Plasmodesmata: the battleground against intruders. Trends in Plant Science 16, 201–210. doi:10.1016/j.tplants.2011.01.004

Lucas, W.J., Bouchepillon, S., Jackson, D.P., Nguyen, L., Baker, L., Ding, B., and Hake, S. (1995). Selective Trafficking of Knotted1 Homeodomain Protein and Its mRNA through Plasmodesmata. Science 270, 1980–1983. doi:10.1126/science.270.5244.1980

Lucas, W.J., Ham, L.K., and Kim, J.Y. (2009). Plasmodesmata-bridging the gap between neighboring plant cells. Trends in Cell Biology 19, 495–503. doi:10.1016/j.tcb.2009.07.003

Magyar, Z., De Veylder, L., Atanassova, A., Bakó, L., Inzé, D., and Bögre, L. (2005). The role of the Arabidopsis E2FB transcription factor in regulating auxin-dependent cell division. Plant Cell 17, 2527–41. doi: 10.1105/tpc.105.033761

Maule, A.J. (2008). Plasmodesmata: structure, function and biogenesis. Current Opinion in Plant Biology 11, 680–686. doi:10.1016/j.pbi.2008.08.002

Miyazawa, Y., Nakajima, N., Abe, T., Sakai, A., Fujioka, S., Kawano, S., Kuroiwa, T., and Yoshida, S. (2003). Activation of cell proliferation by brassinolide application in tobacco BY-2 cells: effects of brassinolide on cell multiplication, cell-cycle-related gene expression, and organellar DNA contents. Journal of Experimental Botany 54, 2669–2678. doi:10.1093/jxb/erg312

Nagata, T., Nemoto, Y., and Hasezawa, S. (1992). Tobacco BY-2 cell line as the ‘Hela’ cell in the cell biology of higher plants. International review of cytology 132, 1–30. doi:10.1016/S0074-7696(08)62452-3

Nakajima, K., Sena, G., Nawy, T., and Benfey, P. N. (2001). Intercellular movement of the putative transcription factor SHR in root patterning. Nature 413, 307–11. doi: 10.1038/35095061

Nicolas, W.J., Grison, M.S., Trepout, S., Gaston, A., Fouche, M., Cordelieres, F.P., Oparka, K., Tilsner, J., Brocard, L., and Bayer, E.M. (2017). Architecture and permeability of post-cytokinesis plasmodesmata lacking cytoplasmic sleeves. Nature Plants 3, 17082. doi:10.1038/nplants.2017.82.

Oparka, K. J., Roberts, A. G., Boevink, P., Santa Cruz, S., Roberts, I., Pradel, K. S., et al. (1999). Simple, but not branched, plasmodesmata allow the nonspecific trafficking of proteins in developing tobacco leaves. Cell 97, 743–54. doi: 10.1016/s0092-8674(00)80786-2

Roberts, A.G., and Oparka, K.J. (2003). Plasmodesmata and the control of symplastic transport. Plant Cell and Environment 26, 103–124. doi:10.1046/j.1365-3040.2003.00950.x

Schöffl, F., Raschke, E., and Nagao, R.T. (1984). The DNA sequence analysis of soybean heat-shock genes and identification of possible regulatory promoter elements. EMBO Journal 3, 2491–2497. doi:10.1002/j.1460-2075.1984.tb02161.x

Sessions, A., Yanofsky, M. F., and Weigel, D. (2000). Cell-cell signaling and movement by the floral transcription factors LEAFY and APETALA1. Science 289, 779–82. doi: 10.1126/science.289.5480.779

Stonebloom, S., Burch-Smith, T., Kim, I., Meinke, D., Mindrinos, M., and Zambryski, P. (2009). Loss of the plant DEAD-box protein ISE1 leads to defective mitochondria and increased cell-to-cell transport via plasmodesmata. Proc. Natl. Acad. Sci. U. S. A. 106, 17229–34. doi: 10.1073/pnas.0909229106

Turgeon, R., and Wolf, S. (2009). Phloem Transport: Cellular Pathways and Molecular Trafficking. Annual Review of Plant Biology 60, 207–221. doi:10.1146/annurev.arplant.043008.092045

Wu, X., Dinneny, J. R., Crawford, K. M., Rhee, Y., Citovsky, V., Zambryski, P. C., et al. (2003). Modes of intercellular transcription factor movement in the Arabidopsis apex. Development 130, 3735–45. doi: 10.1242/dev.00577

Xu, M., Cho, E., Burch-Smith, T. M., and Zambryski, P. C. (2012). Plasmodesmata formation and cell-to-cell transport are reduced in decreased size exclusion limit 1 during embryogenesis in Arabidopsis. Proc. Natl. Acad. Sci. U. S. A. 109, 5098–103. doi: 10.1073/pnas.1202919109

Xu, X.M., and Jackson, D. (2010). Lights at the end of the tunnel: new views of plasmodesmal structure and function. Current Opinion in Plant Biology 13, 684–692. doi:10.1016/j.pbi.2010.09.003

Yamaoka, N., Shimizu, K., Imaizumi, Y., Ito, T., Okada, Y., and Honda, H. (2019). Open-Chamber Co-Culture Microdevices for Single-Cell Analysis of Skeletal Muscle Myotubes and Motor Neurons with Neuromuscular Junctions. Biochip Journal 13, 127–132. doi:10.1007/s13206-018-3202-3

