## Supplemental file for "Quantitative Analysis of Plasmodesmata Permeability using Cultured Tobacco BY-2 Cells Entrapped in Microfluidic Chips"

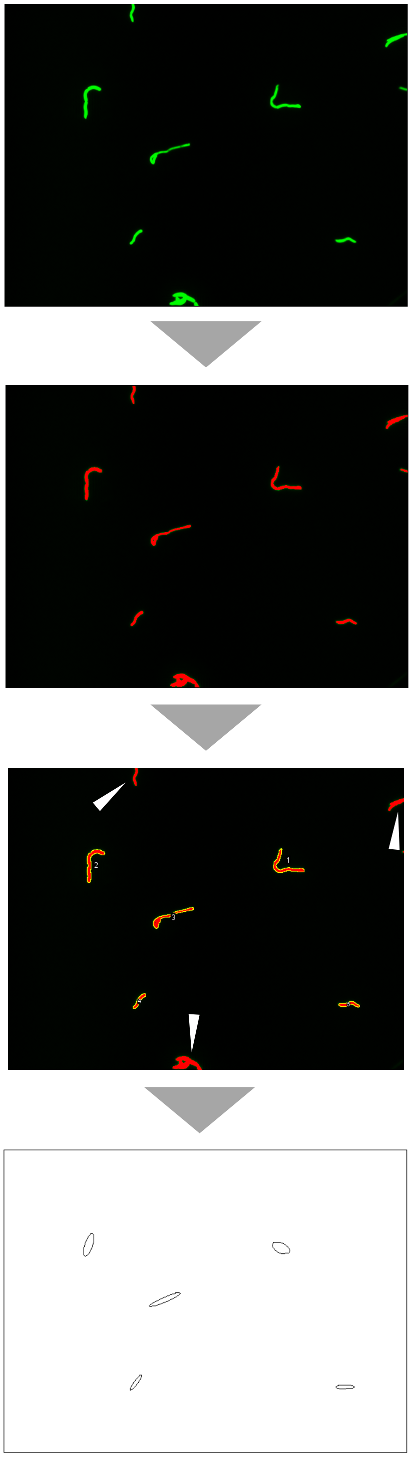

(Step 1) Acquiring of fluorescent images of BY2 cells stained with Celltracker Green.

(Step 2) Recognizing the cells as objects by color threshold of ImageJ.

(Step 3) Excluding the objects crossing the edge of the image (white arrowhead) using ImageJ.

(Step 4) Analyzing the objects with an approximation by ellipses using ImageJ and measuring the length of minor axis of the ellipses and the area.

**Supplementary Figure 1** Procedure for image analysis of BY-2 cells using ImageJ.

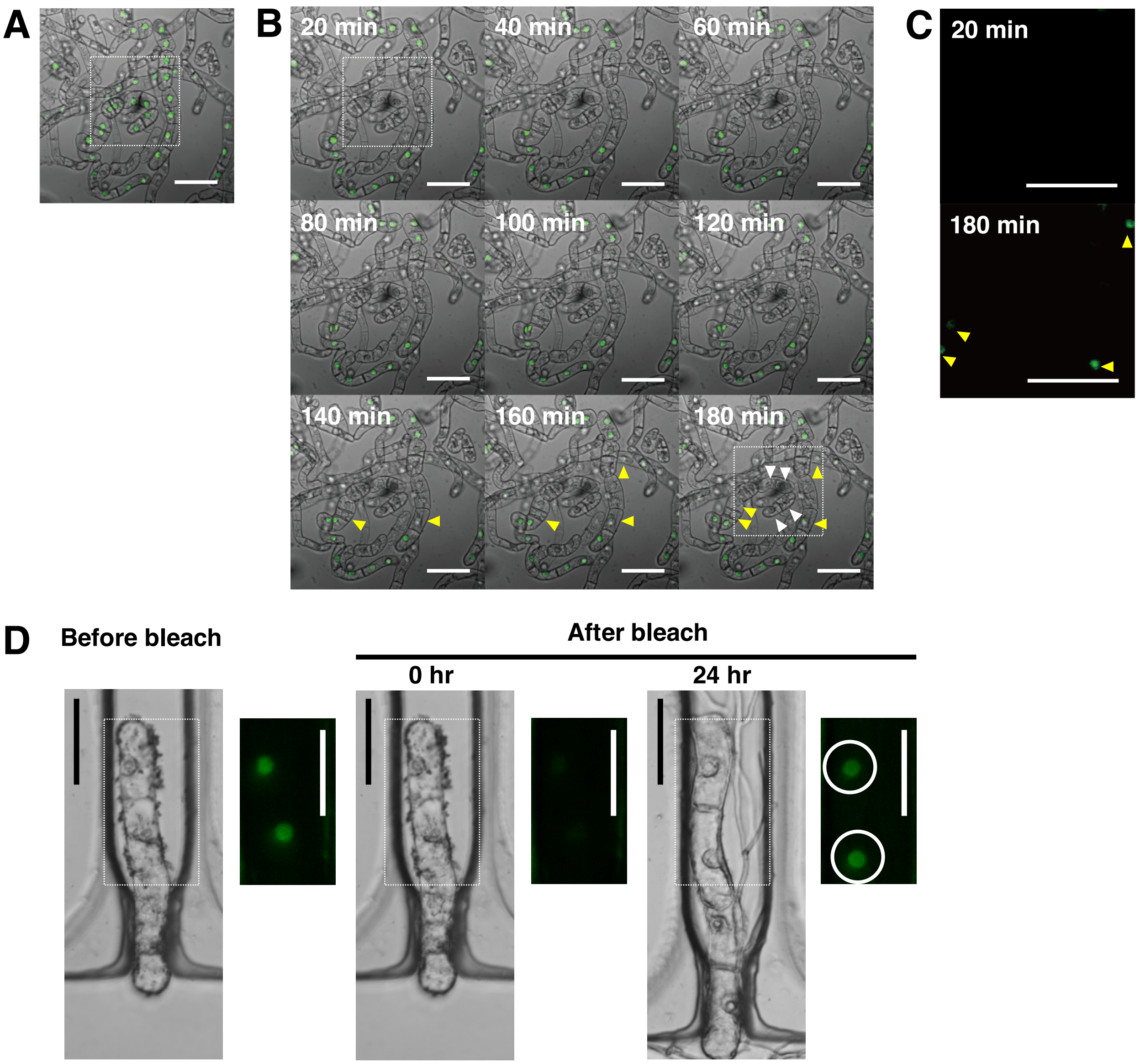


**Supplementary Figure 2** Determination of fluorescence recovery after photobleaching (FRAP) experimental conditions. **(A)** Laser scanning microscope image of BY-2 cells expressing H2B-GFP. Slashed rectangle indicates laser-irradiated area in (B). **(B)** Representative images of FRAP. Yellow and white arrowheads indicate nuclei with recovered and unrecovered GFP fluorescence, respectively. Slashed rectangles indicate areas magnified in (C). **(C)** Magnified fluorescence image of recovered GFP. **(D)** Microscopic images of BY-2 cells in microfluidic device before and after photobleaching. Slashed rectangles indicate areas of fluorescence images. Circles indicate signal from *de novo*-expressed GFP at 24 h after photobleaching. Confocal laser microscopy images were merged from 10 consecutive optical sections. Fluorescence images were merged with bright-field images in (A, B). Scale bars, 200 µm (A–C), 50 µm (D).
